# MAPKAP Kinase 2 Orchestrates Memory T Cell Inflation in Cytomegalovirus Infection

**DOI:** 10.64898/2026.01.16.699974

**Authors:** Eleni Panagioti, Xueyang Yu, Yi Wen Kong, Kristina Makakova, Noe B. Mercado, Sean Edward Lawler, Michael B. Yaffe, Charles H. Cook

## Abstract

Memory T cell inflation is a distinctive immunological phenomenon observed during persistent viral infections, such as Cytomegalovirus (CMV). Unlike conventional memory T cell responses, which contract after infection resolution, a subset of CMV-specific T cells undergoes a progressive and sustained expansion, termed “inflation”, which is thought to be critical for long-term immune surveillance. The molecular mechanisms that govern memory T cell inflation remain incompletely understood, yet they are pivotal for understanding immune persistence and designing strategies against chronic viral infections. In this study, we investigate the role of MAP kinase-activated protein kinase 2 (MK2), a key downstream effector of p38 MAPK signaling, in regulating T cell responses during murine CMV (MCMV) infection. Using MK2 knockout (MK2-KO) mice, we demonstrate that MK2 deficiency alters the dynamics of MCMV-specific CD8⁺ T cell responses without impairing viral control or tissue replication. MK2 deficiency led to a reduction in non-inflationary MCMV-specific CD8⁺ T cells during the acute phase, followed by enhanced expansion of inflationary CD8⁺ T cell subsets during persistence. Furthermore, MK2-KO mice exhibited impaired effector differentiation, as evidenced by decreased expression of the terminal differentiation marker KLRG1 on MCMV-specific CD8⁺ T cells. Collectively, these findings identify MK2 as a pivotal regulator of CD8⁺ T cell magnitude, kinetics, and phenotype during both acute and chronic MCMV infection. By elucidating the role of MK2 in the regulation of memory T cell inflation, this study provides new mechanistic insight into immune regulation with implications for vaccination, chronic infection, and immune aging.

**Graphical abstract:** The graphical abstract was created using BioRender (https://biorender.com/).

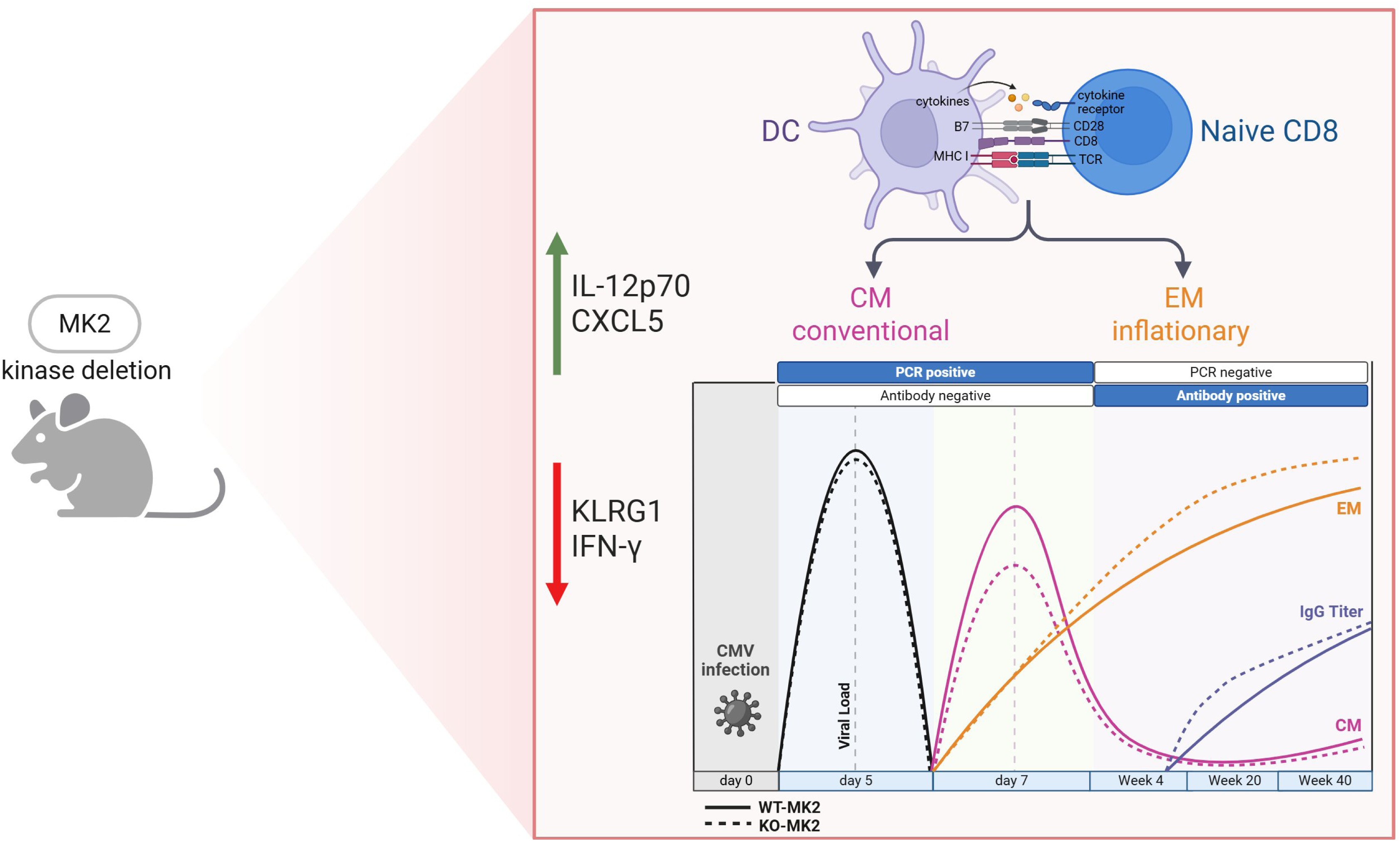

## Introduction

Antiviral T cell responses play a pivotal role in controlling infections. Understanding the molecular mechanisms governing T cell development and differentiation is critical for enhancing these responses^1^. Cytomegalovirus (CMV), a widely prevalent herpesvirus, exhibits seroprevalence rates ranging from 40% to over 90%^2,3^. Although CMV infections are generally asymptomatic in healthy individuals, viral reactivation represents a major clinical challenge in immunocompromised populations^4,5^. Given the widespread prevalence of CMV and the associated health risks, the development of effective therapeutic or preventive CMV vaccines remains a critical public health priority^6,7^.

T cell responses to CMV typically exhibit two distinct patterns of expansion. The conventional pattern is characterized by an initial robust expansion, followed by contraction, and maintenance at low, stable levels. In contrast, a subset of T cells demonstrates an unconventional pattern of persistent expansion, referred to as T cell inflation^8^. These inflationary T cells can remain elevated throughout the host’s lifetime, sustaining a potent CD8^+^ T cell response. They exhibit an effector-memory-like phenotype, retain cytolytic activity, and rapidly produce cytokines such as IFN-γ and TNF-α. Their ability to maintain continuous immune pressure is critical for long-term CMV control, effectively preventing reactivation. Unlike T cells in other chronic infections, inflationary T cells are resistant to functional exhaustion, emphasizing their distinctive role in immune surveillance^8,9^. Elucidating the molecular mechanisms that drive memory T cell inflation may reveal new therapeutic targets for chronic viral infections and inform design and development of next-generation T cell-based therapies.

Accumulating evidence has identified several key factors that influence T cell inflation during persistent viral infections. First, strong and sustained exposure to viral antigens is essential for T cell activation and proliferation, with TCR affinity for CMV-derived peptides directly influencing the degree of inflation^10,11^. Second, constitutive proteasome processing of antigenic epitopes in latently infected cells is also required for robust inflationary responses^12^. Third, cytokines such as IL-12, IL-15, and type I interferons promote tissue T cell survival, proliferation, and differentiation, while IL-10 inhibits memory T cell inflation during MCMV infection^13–15^. Fourth, innate immune cells, including dendritic cells and macrophages, are integral in presenting viral antigens and delivering essential costimulatory signals^16,17^. Fifth, immune checkpoint molecules, including PD-1 and CTLA-4, further regulate T cell responses, balancing activation to prevent overstimulation and functional exhaustion of inflated T cells^18–20^. The balance between effector and memory T cell differentiation is crucial, as memory T cells are better equipped for long-term persistence, whereas effector T cells may exhibit heightened activation but reduced longevity^21,22^. These factors reveal a complex regulatory network governing T cell inflation with ongoing research focused on identifying key signaling pathways that impact the magnitude and functionality of inflationary T cells^23^.

MAPK-Activated Protein Kinase 2 (MK2), a key component of the p38 MAPK signaling pathway, plays a crucial role in regulating inflammatory and stress responses^24–26^. While its role in T cell development remains underexplored, emerging evidence suggests that MK2 may influence T cell function by regulating the production of pro-inflammatory cytokines such as TNF-α and IL-6, which are essential for T cell activation and differentiation^27–29^. MK2 also contributes to T cell proliferation, survival and stress responses, potentially affecting the maintenance of memory T cells in chronic infections and cancer^30–32^. Although MK2 has not been directly implicated in memory T cell inflation, its established role in regulating inflammatory and stress response pathways suggests it may influence T cell survival and immune function.

In this study, we investigated the role of MK2 signaling in modulating T cell inflation during MCMV infection. Our results demonstrate that MK2 knockout (MK2-KO) mice tolerate the absence of MK2 without significant effects on viral replication. However, these mice exhibited marked alterations in MCMV-specific T cell responses. Specifically, MK2-KO mice had fewer non-inflationary MCMV-antigen-specific CD8^+^ T cells, but a notable expansion of inflationary MCMV-antigen-specific CD8^+^ T cells. MK2 signaling was also crucial for the differentiation of effector and memory T cells, as indicated by reduced expression of Killer Cell Lectin-Like Receptor G1 (KLRG1), a marker of terminal effector differentiation. Furthermore, systemic IFN-γ levels were significantly lower during the innate immune response, underscoring MK2’s critical role in shaping T cell magnitude, kinetics, and phenotype during both the acute and chronic phases of MCMV infection.

## Materials and methods

### Mice

Female and male C57BL/6J mice (Strain no. 00664; JAX), aged 7-8 weeks, were purchased from Jackson Laboratory (Bar Harbor, ME, USA). Female and male MK2 knockout (MK2-KO) and MK2 Floxed mice, on a C57BL/6J background were engineered as previously described^33^ and bred at the Koch Institute, Massachusetts Institute of Technology (MIT). Genotyping was performed to confirm MK2 deletion. All mice were maintained under specific-pathogen-free conditions at the Koch Institute and Beth Israel Deaconess Medical Center animal facilities and were 10-14 weeks old at the start of each experiment. All animal procedures adhered to the Institutional Animal Care and Use Committee (IACUC) guidelines of MIT and Beth Israel Deaconess Medical Center.

### Virus Propagation, Infections, and Determination of Viral Load

Recombinant MCMV lacking the m157 gene^34^ (MCMVΔm157) was kindly provided by Ulrich Koszinowski (Ludwig-Maximilians-Universität, Munich, Germany). Virus propagation was performed in NIH-BALB/3T3 clone A31 mouse embryo fibroblasts (ATCC, CCL-163) as previously described^35^. Cells were maintained in Dulbecco’s Modified Eagle’s Medium (DMEM, ATCC, 30-2002) supplemented with 10% heat-inactivated calf bovine serum (ATCC, 30-2030), 100 IU/mL penicillin/100 µg/mL streptomycin (Gibco, 15140112), in a 5% CO2 environment at 37°C. Passaging was performed every 3-4 days, and trypsin-EDTA (Gibco, 25200056) was removed by centrifugation. Viral titers were determined by plaque assays using NIH-BALB/3T3 cells^35^. Age- and gender-matched MK2-KO and MK2 wildtype mice were infected intraperitoneally (i.p.) with 5 × 10⁴ plaque-forming units (PFU) of MCMV.

Genomic DNA was extracted from snap-frozen mouse tissues (brain, lungs, spleen, liver, and salivary glands) collected at days 6 and 14 post-infection using the DNeasy Blood & Tissue Kit (Qiagen, Cat no. 69504). Viral load was quantified by droplet digital PCR (ddPCR) targeting the MCMV glycoprotein B (gB) gene using the QX200 ddPCR System (Bio-Rad) and TaqMan Master Mix (Thermo Fisher Scientific), following the manufacturers’ protocols. Aliquots (100 ng) of DNA were used as templates for each reaction. MCMV gB was used to determine viral load, and MCMV copy numbers were calculated using the HindIII plasmid. Data are expressed as the ratio of MCMV per Housekeeping (HK) gene. The following primers were used: MCMV gB (forward: 5′-GAAGATCCGCATGTCCTTCAG-3′, reverse: 5′-AATCCGTCCAACATCTTGTCG-3′) and β-actin (forward: 5′-GATGTCACGCACGATTTCC-3′, reverse: 5′-GGGCTATGCTCTCCCTCAC-3′). ddPCR data were analyzed using Bio-Rad QuantaSoft software.

### MCMV-Specific Antibody Detection

Fresh blood was collected from the tail vein of infected mice at days 30, 100, 140, and 270 post-infection using Microvette® CB 300 Serum capillary collection tubes (SARSTEDT, Cat no. 16.440.100) and allowed to clot for 2 hours at 4°C. After centrifugation at 2000 rpm for 15 minutes, sera were transferred to new tubes and stored at −20°C. MCMV-specific serum IgG levels were quantified using an enzyme-linked immunosorbent assay (ELISA). Briefly, 96-well plates (BioLegend, Cat no. 423501) were coated overnight at 4°C with NIH-BALB/3T3-derived MCMV-Delta157 in bicarbonate buffer (pH 9.6). Plates were blocked for 1 hour at 37°C with PBS containing 5% milk powder. Sera were diluted 1:500 in PBS with 1% milk powder and incubated in the blocked wells for 1 hour at 37°C. Horseradish peroxidase (HRP)-conjugated IgG antibody (Southern Biotech, Birmingham, USA) was diluted 1:4000 in PBS with 1% milk powder and added to the wells for 1 hour at 37°C. For development, 100 μL of TMB substrate (ThermoFisher Scientific, Cat no. 34021) was added to each well and incubated for 15 minutes at room temperature. The reaction was stopped by adding 100 μL of ELISA stop solution (Invitrogen, Cat no. SS04), and absorbance was measured at 450 nm within 5 minutes using a microplate reader (Model 680, Bio-Rad).

### Preparation of Single-Cell Suspensions from Spleen

Mice were lethally anesthetized by carbon dioxide inhalation. Spleens were collected in RPMI-1640 medium supplemented with 10% heat-inactivated FBS (Thermo Fisher Scientific, Cat no. A5670801), 1% Penicillin-Streptomycin (Gibco, Fisher Scientific, Cat no. 15-140-122), and 1% HEPES (Thermo Scientific, Cat no. J61275.AK). Individual spleens were minced, passed through a 70 μm cell strainer (Falcon, Cat no. 08-771-2), erythrocytes were lysed using 1x RBC Lysis buffer (eBioscience, Cat no. 00-4333-57), washed twice with medium, and immunostaining was performed.

### Immune responses by Spatial Flow Cytometry

For immunophenotyping, peripheral blood mononuclear cells (30 µL blood/well) and spleenocytes (1.5 × 10^6 cells/well), were plated in a non-tissue culture-treated 96-well U-bottom plate (Corning, Cat no. 351177). Determination of the MCMV antigen-specific T cell response by MHC class I tetramers was performed as described^36^. MHC class I tetramers specific for the following MCMV epitopes: m45_985–993_, m57_816–824_, m139_419-426_, m38_316–323_, and IE3_416–423_ in C57BL/6 mice^37^ were used. Cells were labeled with Viability Ghost Red APC/Cy7 (TONBO Biosciences, 13-0865, 1:800) or 7-Aminoactinomycin D 7AAD (Thermo Fisher Scientific, A1310, 1:800) in staining buffer (2% FBS, 0.05% sodium azide in PBS) for 15 minutes at room temperature. Cells were then blocked with Fc block (BD, clone 2.4G2, 553142, 1:100) for 10 minutes at 4°C. Following blocking, cells were stained with a mixture of fluorochrome-conjugated antibodies.

The following antibodies were used for surface staining: CD45.2 APC-Cy7 (BioLegend, clone 104, catalog 109823, 1:200), CD8a Violet Fluor 450 (BioLegend; clone 2.43; catalog 75-1886-U100; 1:100), CD8a FITC (BioLegend, clone 53-6.7, catalog 100706, 1:150), PD1 BV605 (BioLegend, clone RMP1-30, catalog 748267, 1:200), CD3 BV711 (BioLegend, clone 17A2, catalog 100241, 1:200), CD3 BV421 (BioLegend, clone 145-2C11, catalog 562600, 1:100), CD44 BV786 (BioLegend, clone IM7, catalog 563736, 1:200), CD44 APC-Cy7 (BioLegend, clone IM7, catalog 560568, 1:200), CD4 Spark Blue 515 (BioLegend, clone GK1.5, catalog 100493, 1:100), NK1.1 Pe/Cy7 (eBioscience, clone PK136, catalog 25-5941-81, 1:200), F4/80 Pe (TONDO biosciences, clone BM8.1, catalog 50-4801, 1:200), Tim3 PerCP/Cy5.5 (BioLegend, clone B8.2C12, catalog 134011, 1:100), KLRG1 FITC (BioLegend, clone 2F1/GKRG1, catalog 138410, 1:100), KLRG1 Brilliant Violet 711 (BioLegend, clone MAFA, 2F1-Ag, catalog 138427, 1:200), CD19 Pe-CF594 (BioLegend, clone 6D5, catalog 115554, 1:200), Gr1 A700 (BioLegend, clone RB6-8C5, catalog 108422, 1:800), CD62L BV650 (BioLegend, clone MEL-14, catalog 564108, 1:200), CD11b Brillian Violet 650 (BioLegend, clone M1/70, catalog 101239, 1:150), and CD11b BV510 (BioLegend, clone M1/70, catalog 101245, 1:200). After antibody staining, cells were washed twice with staining buffer and resuspended in 100 µL of fixation buffer (BD Biosciences) for 10 minutes at 4°C. Following fixation, cells were washed twice with staining buffer. Samples were resuspended in 120 μl staining buffer and analyzed using a Cytec Aurora flow cytometer (Cytec Biosciences). FlowJo software (Treestar Inc. ver 10.) was used for data analysis. The detailed gating strategy is provided in Supplementary Fig. 1.

### Cytokine measurements

For measurement of cytokine and chemokine panels in mice, serum samples were collected at the time points indicated in the figure legends. Cytokine analysis was performed using the ELISA MAX™ Deluxe Set Mouse IL-10 assay kit (BioLegend, Cat no. 431414) and the Luminex™ 200 system (Luminex, Austin, TX, USA) at Eve Technologies Corp. (Calgary, Alberta, Canada). A total of 44 markers were measured simultaneously using Eve Technologies’ Mouse Cytokine 44-Plex Discovery Assay®, which combines a 32-plex and a 12-plex kit (MilliporeSigma, Burlington, Massachusetts, USA). The 32-plex panel included the following cytokines and chemokines: Eotaxin, G-CSF, GM-CSF, IFN-γ, IL-1α, IL-1β, IL-2, IL-3, IL-4, IL-5, IL-6, IL-7, IL-9, IL-10, IL-12(p40), IL-12(p70), IL-13, IL-15, IL-17, IP-10, KC, LIF, LIX, MCP-1, M-CSF, MIG, MIP-1α, MIP-1β, MIP-2, RANTES, TNF-α, and VEGF. The 12-plex panel included 6Ckine/Exodus2, Erythropoietin, Fractalkine, IFN-β1, IL-11, IL-16, IL-20, MCP-5, MDC, MIP-3α, MIP-3β, and TARC. The assay sensitivities for the 44-plex ranged from 0.3 to 30.6 pg/mL, with specific sensitivity values for each analyte provided in the MilliporeSigma MILLIPLEX® MAP protocol.

### Statistical Analyses

Statistical analyses were performed using GraphPad Prism 6.0 software (GraphPad Software). Differences between two groups were assessed using an unpaired Student’s t-test. For comparisons involving more than two groups, one-way ANOVA followed by Tukey’s or two-way ANOVA followed by Šídák’s models were performed to account for multiple comparisons used. A P-value of < 0.05 was considered statistically significant. Final graphs were formatted using Adobe Illustrator (version CC).

## Results

### MK2 Deficiency Does Not Affect Viral Titers in Lymphoid and Non-Lymphoid Organs but Enhances Early Antiviral Antibody Responses

MK2 whole body knockout (MK2-KO) mice generated via conventional gene targeting that were used in these experiments were validated by genotyping prior to experimental analyses^33,38^. Global loss of MK2 does not impact overall survival or developmental parameters, including body weight (Supplementary Fig. 2A). However, MK2-KO mice exhibit reduced total splenocyte counts and a trend toward decreased spleen weight (Supplementary Fig. 2B-C), despite no significant change in absolute numbers of CD3⁺ T cells (Supplementary Fig. 2D). Peripheral blood analyses reveal no differences in frequencies of circulating myeloid populations (Supplementary Fig. 3A), and while untreated MK2-KO mice show increased frequencies of total circulating CD3^+^ cells, proportions of CD4^+^ and CD8^+^ T cells remain unchanged (Supplementary Fig. 3B-D). Notably, CD8⁺ T cells from MK2-deficient mice exhibit reduced expression of KLRG1 and display a shift toward a naïve phenotype, characterized by increased frequencies of CD44^-^ CD62L^+^ cells, and a contraction of effector memory subsets (CD44^+^ CD62L^-^) in both CD4⁺ and CD8⁺ T cell compartments (Supplementary Fig. 3C-D). Frequencies of NK cells are comparable between genotypes, while MK2-KO mice show reduced CD19^+^ B cells (Supplementary Fig. 3E-F).

Following systemic infection with Δm157-MCMV (1×10^4^ pfu) via intraperitoneal injection, MK2-KO mice of both sexes exhibit no overt signs of acute infection or immunopathology and tolerate infection similarly to wild-type controls. To evaluate viral replication dynamics, MCMV titers were quantified in both lymphoid and non-lymphoid tissues at defined peak time points. Viral loads measured on day 6 post-infection (p.i.) in the brain, lungs, spleen, and liver (corresponding to the peak of viral replication in these tissues) and on day 14 p.i. in the brain, lungs, spleen, and salivary glands (when viral replication peaks in this organ) were comparable between MK2-KO and WT mice, suggesting that MK2 is dispensable for early control of viral replication and systemic dissemination (Fig. 1A). To determine whether MK2 influences humoral immunity, serum MCMV-specific IgG titers were measured longitudinally from day 30 to day 270 p.i. (days 30, 100, 145 and 270). Both MK2-KO and WT mice demonstrate progressive accumulation of IgG, consistent with MCMV-specific antibody inflation. However, MK2-deficient mice show significantly elevated IgG levels as early as day 30 post-infection, a difference that persists through day 100. Notably, IgG levels began to decline after day 100 in MK2-KO mice and after day 145 in WT controls, indicating a transition to a later immunological phase (Fig. 1B). These findings suggest that persistent MCMV antigen exposure drives IgG inflation at early times after infection in a manner that is limited in extent by MK2-dependent mechanisms.

**Figure 1:**
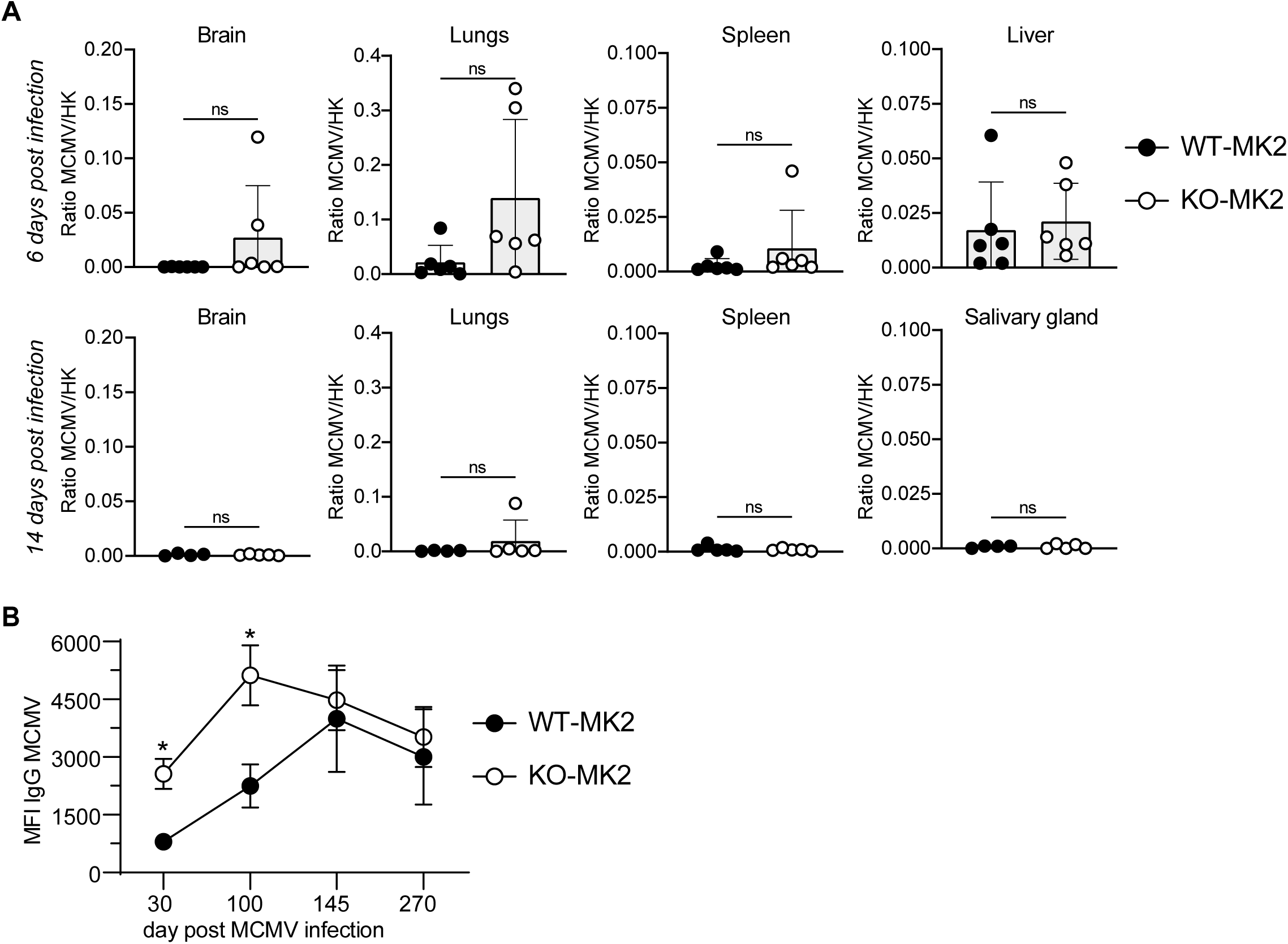
Loss of MK2 Does Not Impact Tissue Viral Burden but Boosts Antiviral IgG Responses. **A.** Viral titers in the brain, lungs, spleen, and liver were quantified on day 6 post-MCMV infection, and in the brain, lungs, spleen, and salivary glands on day 14. Viral load is shown as ratio MCMV/HK values (n = 4-6 mice per group). Statistical comparisons between WT-MK2 and KO-MK2 mice were performed using an unpaired Student’s *t*-test. Data are presented as mean ± SD. Statistical significance: P > 0.05, ns, not significant. **B.** Longitudinal analysis of MCMV-specific IgG titers in WT-MK2 and KO-MK2 mice at days 30, 100, 145, and 270 post-infection. Median fluorescence intensity (MFI) of MCMV-specific IgG is shown (n = 4-6 mice per group). Statistical analysis was performed using two-way ANOVA followed by Šídák’s multiple comparisons test. Data are presented as mean ± SD. Statistical significance: *, *P* < 0.05; P > 0.05, ns, not significant.

### Global MK2 Ablation Influences Magnitude and Kinetics of MCMV-Specific CD8⁺ T Cell Responses

To investigate the role of MK2 in shaping virus-specific CD8⁺ T cell immunity, we longitudinally analyzed peripheral blood from MK2-KO and WT C57BL/6 mice following intraperitoneal infection with Δm157-MCMV. We used H-2K^b^ MHC class I tetramers to quantify CD8⁺ T cells specific for both non-inflationary (m45_985-993_ and m57_816-824_) and inflationary immunodominant epitopes (m139_419-426_, m38_316-323_, and IE3_416-423_), which reflect distinct kinetics of antigen presentation and memory T cell maintenance during MCMV infection^39^.

MK2 deficiency significantly alters both the magnitude and temporal dynamics of MCMV CD8⁺ T cell responses. For non-inflationary epitopes, MK2-KO mice exhibit reduced CD8⁺ T cell frequencies at the peak of the primary antiviral response 7 days after infection (Fig. 2A). This difference is transient and normalizes during the contraction and memory phases, suggesting that MK2 supports the optimal expansion of CD8⁺ T cells during the early stages of viral infection. Strikingly, although antigen-specific CD8⁺ T cell frequencies for inflationary epitopes are similar to WT controls at early time points (day 7 post-infection), MK2-KO mice exhibit a progressive and sustained increase in these cells during later infection, consistent with exaggerated memory inflation (Fig. 2B). This phenotype is consistently observed across multiple inflationary epitopes, suggesting that MK2 exerts a broad, epitope-independent role in controlling CD8⁺ T cell accumulation during chronic MCMV infection. Collectively, these findings identify MK2 as a key regulator of antigen-specific CD8⁺ T cell responses during viral infection, functioning as a molecular brake on the memory inflation program.

**Figure 2:**
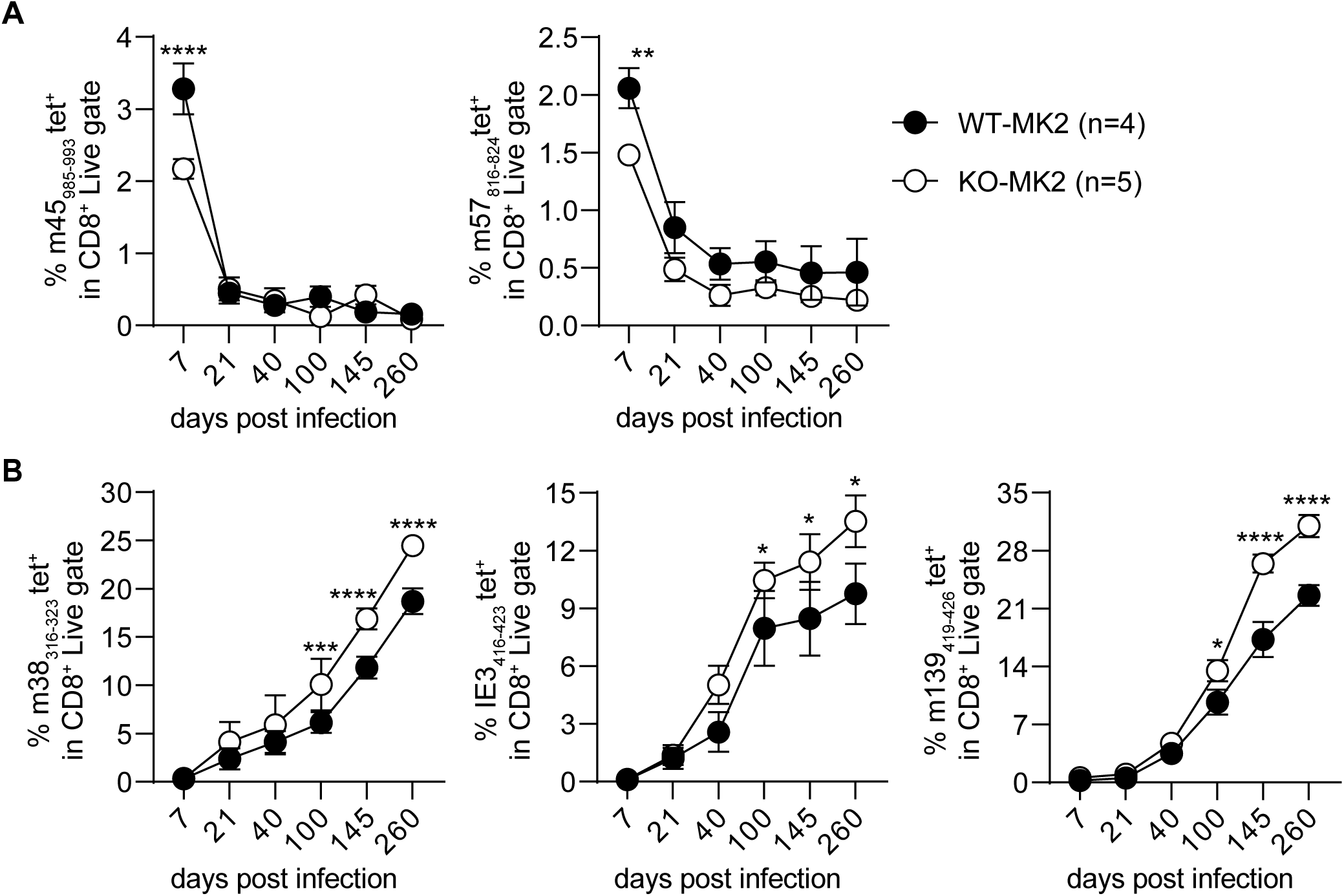
Global MK2 Ablation Alters the Magnitude and Dynamics of MCMV-Specific CD8⁺ T Cell Responses. **A.** Longitudinal analysis of CD8⁺ T cells specific for the non-inflationary MCMV epitopes m45_985-993_ and m57_816-824_ in peripheral blood from WT and KO-MK2 mice (n = 4-8 per group). Statistical comparisons were performed using two-way ANOVA with Šídák’s multiple comparisons test. Data are shown as mean ± SD. Statistical significance: **, *P* < 0.01; ****, *P* < 0.0001; P > 0.05, ns, not significant. Non-significant results are not indicated with a symbol in the graphs. **B.** Longitudinal kinetics of CD8⁺ T cells targeting inflationary MCMV epitopes m38_316-323_, IE3_416-423_ and m139_419-426_ in peripheral blood of WT and KO-MK2 mice (n = 4-8 per group). Statistical analysis was conducted as in **(A)**. Data are presented as mean ± SD. Statistical significance: *, *P* < 0.05; ***, *P* < 0.001; ****, *P* < 0.0001; P > 0.05, ns, not significant. Non-significant differences are not annotated in the graphs.

### MK2 Signaling Controls circulating CD8⁺ T Cell Differentiation and KLRG1 Expression During Chronic MCMV Infection

We next sought to determine how MK2 deficiency affects longitudinal differentiation and functional fitness of MCMV-specific CD8⁺ T cells following systemic ΔMCMV infection. We characterized antigen-specific CD8⁺ T cells in peripheral blood using established memory markers to define effector memory (CD44⁺CD62L⁻), central memory (CD44⁺CD62L⁺), and naïve (CD44⁻CD62L⁺) subsets, along with key differentiation and survival markers, KLRG1 and CD127, respectively. At day 7 post-infection, MK2-KO mice exhibit reduced frequencies of circulating KLRG1⁺CD127⁻ CD8⁺ T cells specific for both inflationary and non-inflationary MCMV epitopes, compared to WT (Fig. 3A). By day 100 post-infection, however, these differences are no longer apparent (Fig. 3B). The distribution of CD8⁺ T cell subsets based on CD44 and CD62L expression is comparable between MK2-KO and WT mice during the acute infection phase (Fig. 3C). In the chronic phase, MK2-KO mice demonstrate increased proportions of effector memory (CD44⁺CD62L⁻) and concomitant decreases in central memory (CD44⁺CD62L⁺) CD8⁺ T cells, particularly for non-inflationary epitopes (Fig. 3D). Notably, longitudinal analysis reveals consistent reductions in KLRG1 expression in MCMV-specific CD8⁺ T cells from MK2-KO mice across all epitopes on day 7 post-infection, irrespective of their inflationary status. This difference, however, disappears by later time points (Fig. 3E). KLRG1 is a well-characterized marker of terminally differentiated effector CD8⁺ T cells, with expression closely linked to the acquisition of cytotoxic functions critical for viral control, particularly within inflationary CD8⁺ T cell populations^40,41^. The marked reduction in KLRG1⁺ cells observed in MK2-KO mice implicates MK2 as a key regulator of CD8⁺ T cell terminal differentiation, influencing the balance between effector and memory-like fates.

**Figure 3:**
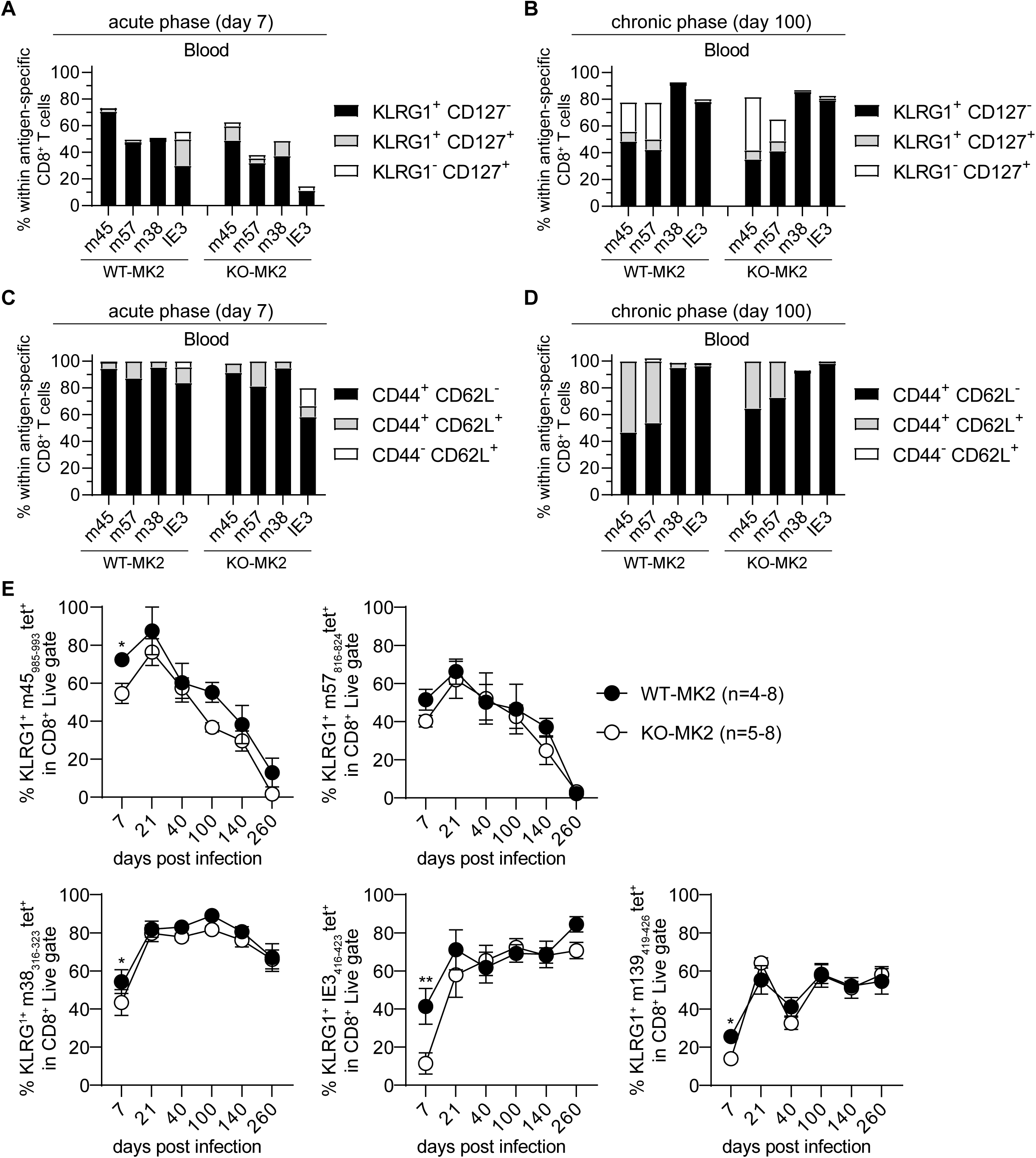
MK2 Signaling Shapes CD8⁺ T Cell Differentiation and KLRG1 Expression During Persistent MCMV Infection. **A.** Frequencies of KLRG1⁺CD127⁻, KLRG1⁺CD127⁺, and KLRG1⁻CD127⁺ MCMV epitope-specific CD8⁺ T cells in peripheral blood of WT and KO-MK2 mice at day 6 post-infection. Data are presented as mean ± SD. **B.** Frequencies of KLRG1⁺CD127⁻, KLRG1⁺CD127⁺, and KLRG1⁻CD127⁺ MCMV epitope-specific CD8⁺ T cells in peripheral blood at day 100 post-infection. Data are presented as mean ± SD. **C.** Proportions of effector memory (CD44⁺CD62L⁻), central memory (CD44⁺CD62L⁺), and naïve (CD44⁻CD62L⁺) MCMV epitope-specific CD8⁺ T cells in peripheral blood of WT and KO-MK2 mice at day 6 post-infection. Data are presented as mean ± SD. **D.** Proportions of effector memory, central memory, and naïve MCMV-specific CD8⁺ T cells at day 100 post-infection. Data are presented as mean ± SD. **E.** Kinetics of KLRG1⁺ CD8⁺ T cells specific for non-inflationary epitopes m45_985-993_ and m57_816-824_ in peripheral blood from WT and KO-MK2 mice at days 7, 21, 40, 100, 140, and 260 post-infection (n = 4-8 per group). Statistical analysis was performed using two-way ANOVA with Šídák’s multiple comparisons test. Data are presented as mean ± SD. Statistical significance: *, *P* < 0.05; P > 0.05, ns, not significant. Non-significant differences are not indicated. **F.** Kinetics of KLRG1⁺ CD8⁺ T cells specific for inflationary epitopes m38_316-323_, IE3_416-423_, and m139_419-426_ in peripheral blood from WT and KO-MK2 mice (n = 4-8 per group). Statistical analysis was performed as in **(E)**. Statistical significance: *, *P* < 0.05; **, *P* < 0.01; P > 0.05, ns, not significant. Non-significant differences are not annotated in the graphs.

Our findings identify MK2 as a previously unrecognized regulator of CD8⁺ T cell differentiation during chronic viral infection, with important implications for antiviral immunity. By modulating KLRG1 expression, MK2 influences the balance between effector and memory-like CD8⁺ T cell fates, revealing a novel layer of regulation in persistent infection.

### MK2 Signaling Regulates the Magnitude and Differentiation of MCMV-Specific CD8⁺ T Cells in the Spleen

After longitudinally assessing CD8⁺ T cell responses in peripheral blood, we next examined the spleen, a key secondary lymphoid organ essential for antiviral immunity. Spleens were harvested from MK2-KO and WT mice following systemic ΔMCMV infection for flow cytometric analysis of antigen-specific CD8⁺ T cell frequency and phenotype. Interestingly, MK2-KO mice exhibit reduced immune responses in the spleen during acute MCMV infection, as evidenced by significantly diminished splenomegaly compared to infected WT controls (Fig. 4A). Spleen weights are lower in infected MK2-KO mice, accompanied by a marked reduction in total splenocyte numbers, including CD3⁺ T cells (Fig. 4B-D). These findings suggest that MK2 contributes to the regulation of splenic inflammation during acute viral infection. To further characterize immune responses, we performed flow cytometry analysis of T cells, myeloid cells, and MHC class I tetramer⁺ CD8⁺ T cells specific for non-inflationary (m45_985-993_, m57_816-824_) and inflationary (m38_316-323_, IE3_416-423_, m139_419-426_) MCMV epitopes during early infection. For CD4^+^ cells MK2-KO mice exhibit reduced absolute numbers of splenic CD4⁺ T cells, including decreases in KLRG1⁺ and CD44^+^CL62L^-^ effector memory CD4^+^ T cell subsets (Fig. 4E).

**Figure 4:**
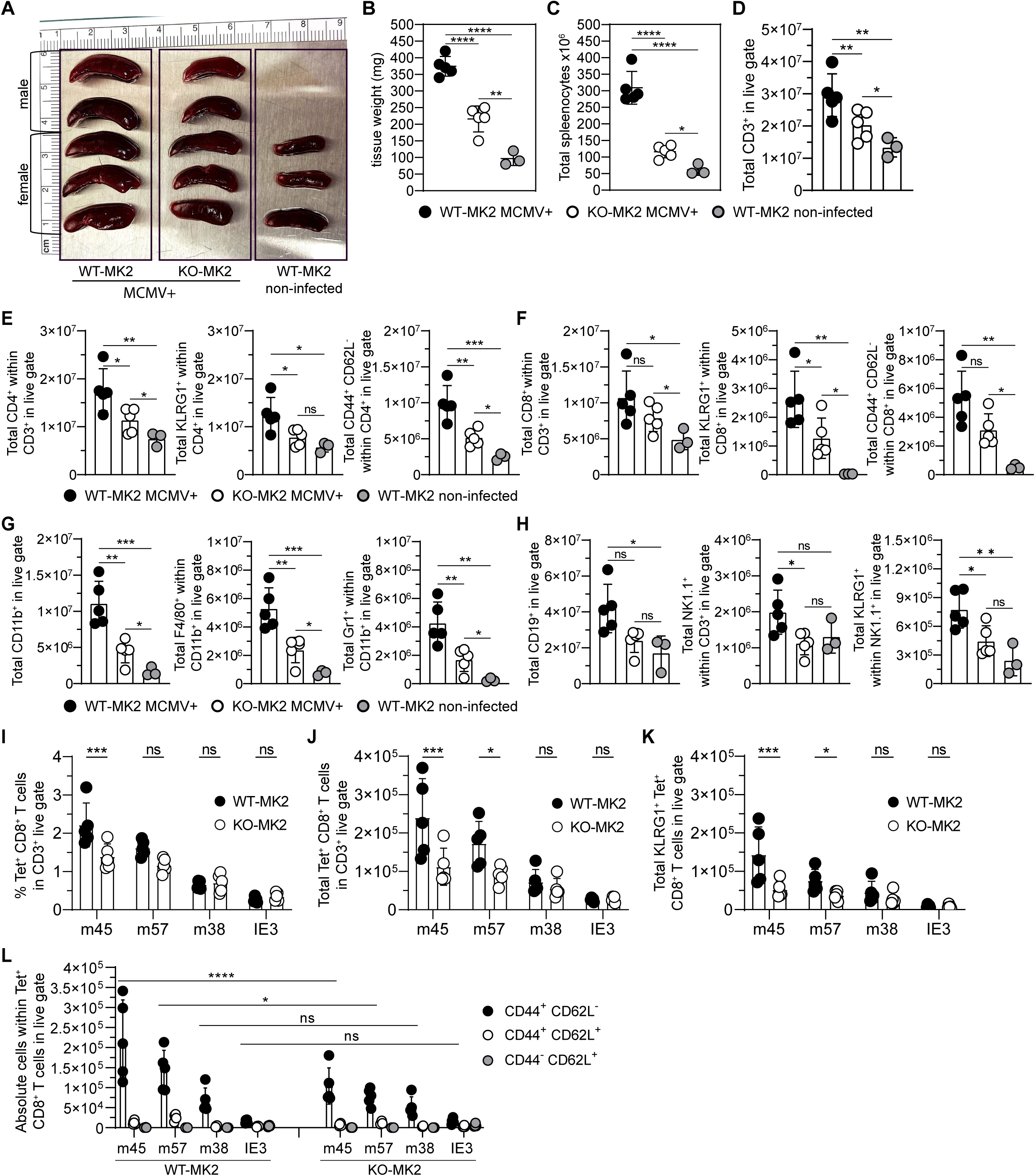
MK2 Signaling Shapes the Magnitude and Effector Differentiation of Splenic CD8⁺ T Cells During MCMV Infection. **A.** Representative images of spleens from WT and KO-MK2 male and female mice at day 7 post MCMV infection. Spleens from non-infected naïve WT mice are shown as controls. WT spleens exhibit marked splenomegaly, with a reduced size increase in MK2-deficient mice. **B.** Spleen weights (mg) from mice in **(A)**. Data are mean ± SD. Statistical analysis by one-way ANOVA with Tukey’s multiple comparisons test. Significance: *, *P* < 0.05; **, *P* < 0.01; ****, *P* < 0.0001; P > 0.05, ns, not significant. Panels **C-F** show absolute cell counts at 7 days post MCMV infection. **C.** Total splenocyte counts in WT vs. KO-MK2 infected mice. Naïve WT controls included. Data are mean ± SD; statistical analysis as in **(B)**. **D.** Total numbers of CD3⁺ T cells in spleens of WT vs. KO-MK2 infected mice. Spleens from non-infected naïve WT mice are shown as controls. Data are mean ± SD; statistical analysis as in **(B)**. **E.** Absolute numbers of total CD4⁺ T cells, KLRG1⁺CD4⁺ T cells, and effector memory (CD44⁺CD62L⁻) CD4⁺ T cells in WT and KO-MK2 spleens. Naïve WT controls included. Data are mean ± SD; statistical analysis as in **(B)**. **F.** Absolute numbers of total CD8⁺ T cells, KLRG1⁺CD8⁺ T cells, and effector memory CD8⁺ T cells in WT and KO-MK2 spleens. Naïve WT controls included. Data are mean ± SD; statistical analysis as in **(B)**. **G.** Absolute numbers of CD11b⁺ myeloid cells, F4/80⁺CD11b⁺ macrophages, and Gr1⁺CD11b⁺ MDSCs in WT and KO-MK2 spleens. Naïve WT controls included. Data are mean ± SD; statistical analysis as in **(B)**. Significance: *, *P* < 0.05; **, *P* < 0.01; ***, *P* < 0.001. **H.** Absolute numbers of CD19⁺ B cells, NK1.1⁺CD3⁻ NK cells, and KLRG1⁺NK1.1⁺ cells in WT and KO-MK2 spleens. Naïve WT controls included. Data are mean ± SD; statistical analysis as in **(B)**. **I.** Percentage of MCMV tetramer⁺ CD8⁺ T cells in spleens of WT and KO-MK2 infected mice. Data are mean ± SD; statistical analysis as in **(B)**. **J.** Absolute numbers of MCMV tetramer⁺ CD8⁺ T cells in spleens of WT and KO-MK2 mice. Data are mean ± SD; statistical analysis as in **(B)**. **K.** Absolute numbers of KLRG1⁺ MCMV tetramer⁺ CD8⁺ T cells in spleens of WT and KO-MK2 mice. Data are mean ± SD; statistical analysis as in **(B)**. **L.** Absolute numbers of effector memory (CD44⁺CD62L⁻), central memory (CD44⁺CD62L⁺), and naïve (CD44⁻CD62L⁺) MCMV tetramer⁺ CD8⁺ T cells in spleens of WT and KO-MK2 mice. Statistical comparisons between WT and KO-MK2 groups per epitope were performed using one-way ANOVA with Tukey’s multiple comparisons test. Data are mean ± SD; significance: *, *P* < 0.05; ****, *P* < 0.0001; P > 0.05, ns, not significant.

Total CD8⁺ T cell numbers are also lower in infected MK2-KO mice, along with reduction in KLRG1-expressing CD8⁺ T-cells (Fig. 4F). Within the myeloid compartment, MK2 deficiency results in diminished frequencies of myeloid populations, including macrophages and myeloid-derived suppressor cells (MDSCs), indicating substantial impairment in the antiviral inflammatory myeloid response during acute infection (Fig. 4G). Similarly, analysis of other immune lineages reveals reduced numbers of CD19⁺ B cells, NK cells, and decreased KLRG1 expression on NK cells in infected MK2-KO mice (Fig. 4H).

Analysis of epitope-specific CD8⁺ T-cells during early infection reveals selective impacts of MK2 deficiency on non-inflationary MCMV responses. MK2-KO mice showed reduced frequencies of m45_985-993_-specific CD8⁺ T cells, while the frequencies of CD8⁺ T cells targeting inflationary epitopes remained largely unaffected (Fig. 4I). Absolute cell counts further confirmed that CD8⁺ T cell responses are significantly diminished for non-inflationary epitopes in MK2-KO mice, with no significant differences observed for inflationary epitopes (Fig. 4J). Similarly, frequencies of KLRG1⁺ CD8⁺ T cells specific for non-inflationary epitopes are reduced in spleens of MK2-KO mice, whereas those targeting inflationary epitopes remain unaffected (Fig. 4K). Phenotypic analysis reveals decreased proportions of CD44⁺CD62L⁻ effector CD8⁺ T cells specific for non-inflationary epitopes (m45_985-993_ and m57_816-824_) in MK2-KO infected mice, while inflationary epitope-specific populations showed minimal differences (Fig. 4L).

Together, these findings indicate that MK2 modulates the magnitude and differentiation of CD8⁺ T cells responses to non-inflationary MCMV epitopes during acute infection. While MK2 loss disrupts multiple components of the antiviral immune response, including myeloid, T cell, NK, and B cell compartments, the inflationary CD8⁺ T cell pool remains largely intact, likely maintained through persistent antigen presentation or compensatory pathways active during early infection.

### MK2 Regulates Systemic Cytokine Production During MCMV Infection

MK2 functions as a critical post-transcriptional regulator of key proinflammatory cytokines, including TNF-α, IL-6, and IL-1β, primarily by modulating mRNA stability downstream of p38 MAPK signaling^42^. Because MK2 is central to inflammatory responses, and seems to perturb normal immune responses to MCMV, we investigated MK2’s effect on systemic cytokine secretion during MCMV infection. We thus longitudinally profiled circulating cytokines in MK2-KO and WT mice from baseline (day −14) through days 3 and 7 post-infection. Serum samples were analyzed using multiplex panels covering a broad spectrum of cytokines involved in both innate and adaptive immunity, including Eotaxin, G-CSF, GM-CSF, IFN-γ, IL-1α, IL-1β, IL-2, IL-3, IL-4, IL-5, IL-6, IL-7, IL-9, IL-10, IL-12 (p40), IL-12 (p70), IL-13, IL-15, IL-17, IP-10, KC, LIF, LIX, MCP-1, M-CSF, MIG, MIP-1α, MIP-1β, MIP-2, RANTES, TNF-α, and VEGF as well as the less common 6Ckine/Exodus2, erythropoietin, fractalkine, IFN-β1, IL-11, IL-16, IL-20, MCP-5, MDC, MIP-3α, MIP-3β, and TARC.

Notably, IL-12p70 and CXCL5 are significantly elevated in MK2-KO mice at baseline, and remain high following infection (Fig. 5A). As expected, MCMV infection induces robust increases in serum IFN-γ levels by day 3 in WT mice, whereas this response is markedly blunted for MK2-KO mice (Fig. 5B). In contrast, MK2-KO mice display pronounced elevation in IL-10 levels during very early infection, while WT mice exhibit comparatively muted IL-10 responses (Fig. 5B). Concentrations of other cytokines without significant differences between groups are shown in Supplementary Fig. 4. These results suggest that MK2 plays a crucial role in orchestrating early innate antiviral responses, potentially by modulating the magnitude and duration of cytokine signaling necessary for effective T cell activation and programming.

**Figure 5:**
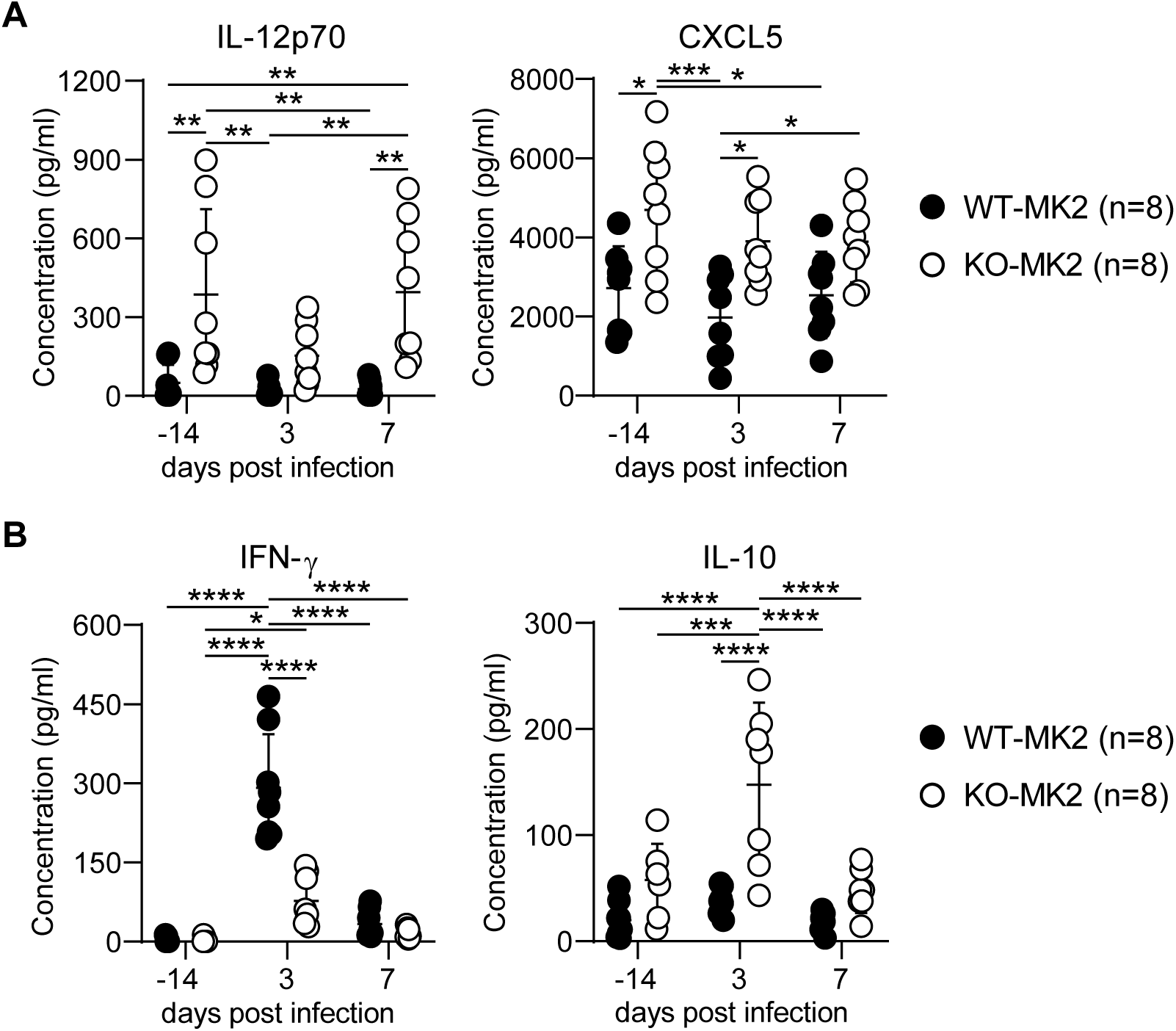
MK2 Regulates Systemic Inflammatory Cytokine Responses During Acute MCMV Infection. **A.** Serum levels of pro-inflammatory cytokines IL-12p70 and CXCL5 in WT and KO-MK2 mice at baseline (day −14) and at days 3 and 7 post-MCMV infection. Cytokine concentrations are reported in pg/mL. Each point represents an individual mouse (n = 8 mice per group). All samples were run in duplicate, and the mean of replicates is shown. Data for the entire group are presented as mean ± SD. Statistical comparisons were performed using two-way ANOVA followed by Tukey’s multiple comparisons test. Significance: *, *P* < 0.05; **, *P* < 0.01; ***, *P* < 0.001; P > 0.05, ns, not significant. **B.** Serum IFN-γ and IL-10 concentrations at the indicated timepoints pre- and post-infection. Data are presented as mean ± SD.; statistical analysis as in **(A)**. Significance: *, *P* < 0.05; ****, *P* < 0.0001; P > 0.05, ns, not significant.

## Discussion

CD8 T cell responses during chronic herpes-viral infections are thought to be essential for maintaining durable immune surveillance and thereby viral control. In this study, we investigate the contribution of MK2 to antiviral immunity after MCMV infection. Our data demonstrate that MK2 deficiency does not impair viral control but significantly reshapes the equilibrium between inflationary and non-inflationary T cell subsets, underscoring a key role for MK2 in directing T cell-mediated immune responses. Specifically, MK2-KO mice exhibit marked impairment in early non-inflationary MCMV-specific CD8⁺ T cells specific for traditional central memory epitopes m45_985-993_ and m57_816-824_, followed by enhanced expansion of inflationary CD8⁺ T cells recognizing m38_316-323_, IE3_416-423_ and m139_419-426_ epitopes.

In our model, MK2 deficiency is associated with reduced expression of KLRG1, a critical marker of terminal effector differentiation, highlighting the essential role of MK2 signaling in orchestrating both effector and memory CD8⁺ T cell maturation. Notably, Baumann et al. previously demonstrated that the magnitude of the inflationary T cell pool during latency is dictated by the abundance of early-primed KLRG1⁻ cells generated during the acute infection, rather than by constraints within peripheral tissue niches^10,43^. These findings align with earlier reports indicating that the maintenance of inflationary T cells is, at least in part, sustained by T cells primed early in infection^10^. Consistent with this model, MK2-KO mice display an expanded pool of KLRG1⁻ CD8⁺ T cells, which is followed by augmented inflationary CD8⁺ T cell responses. This observation is particularly striking given the limited understanding of the molecular pathways controlling KLRG1 surface expression. Collectively, these findings identify MK2 as a key regulator of CD8⁺ T cell differentiation and memory inflation.

IL-10 has previously been identified as a negative regulator of memory T cell inflation^15^. In our model, MCMV-infected MK2-deficient mice exhibit a transient increase in serum IL-10 levels early after infection (day 3), which returns to baseline by day 7, with no significant differences between genotypes at later time points. This supports a role for MK2 in modulating early IL-10 production. IL-12p70 levels remained consistently elevated across time points in MK2-KO. In contrast, IL-2 and IL-15 cytokines, previously implicated in the regulation of memory inflation^9,13,44–46^ were not significantly different between knockout and wild type mice. Notably, antigen-specific responses to non-inflationary m45_985-993_ and m57_816-824_ MCMV epitopes were reduced at day 7 post-infection in MK2-deficient mice, suggesting that early IL-10 elevation may impact the priming of these responses. However, memory inflation to epitopes m38_316-323_, IE3_416-423_ and m139_419-426_ were enhanced at later time points, indicating that early effects of low IL-10 in MK2-KO were likely transient and primarily impacted early priming rather than long-term inflation. Curiously, enhanced memory CD8 T cell inflation in knockout mice occurred despite significantly lower serum IFN-γ at day 3. Taken together, these data suggest that the abundance of early-primed CD8⁺ T cells, particularly the KLRG1⁻ subset during the priming phase, together with enhanced IL-12p70 and CXCL5 signaling, may play a role as important as the previously described influences of IL-2, IL-10, IL-15 and IFN-γ in determining the ultimate magnitude of the inflationary T cell pool during MCMV latency.

The expanded inflationary CD8⁺ T cell compartment observed in MK2-deficient mice likely contributes to sustained antiviral pressure and prolonged CMV control. However, this shift in T cell dynamics underscores the importance of MK2 in maintaining immune balance, restraining excessive inflammation that could compromise T cell persistence or promote immunopathology. These findings suggest that MK2-deficient T cells may have utility in therapeutic settings where robust effector responses are beneficial, such as vaccination or tumor immunotherapy. Nonetheless, the potential risks of immune overactivation warrant careful evaluation. Future studies in controlled vaccination models will be essential to assess the safety and translational potential of MK2-targeted approaches. Moreover, genetic variation in MK2 may influence individual susceptibility to viral pathogenesis, responsiveness to vaccination, and the quality of immune responses in both infectious diseases and cancer.

In summary, MK2 emerges as a key regulator of inflammation, CD8⁺ T cell differentiation, and the mechanisms by which T-cells ultimately control latent CMV infection and effector-memory T-cell inflation responses during MCMV infection. Further studies are warranted to elucidate its broader role in antiviral immunity and to explore the therapeutic potential of targeting MK2 in chronic viral infections and cancer.

## Acknowledgments

We thank the Flow Cytometry Core Facility at BIDMC for their technical assistance. We thank the NIH Tetramer Core Facility (NIH Contract 75N93020D00005 and RRID: SCR_026557) for providing tetramers.

## Author contributions (CRediT details)

E.P. designed the experimental approach, performed the experiments, analyzed and interpreted the data, and wrote the manuscript. X.Y and Y.W.K generated and interpreted data, and contributed to manuscript writing. M.K. performed experiments, analyzed and interpreted data, and contributed to manuscript writing. N.B.M provided viral reagents and contributed to manuscript writing. S.E.L., M.B.Y., and C.H.C. designed the experimental approach, interpreted the data, and contributed to manuscript writing.

## Funding

This work was supported by NIH grants R01CA263324 (to S.E.L and C.H.C.).

## Conflicts of Interest

The authors declare no competing interests.

---

**Figure S1.**
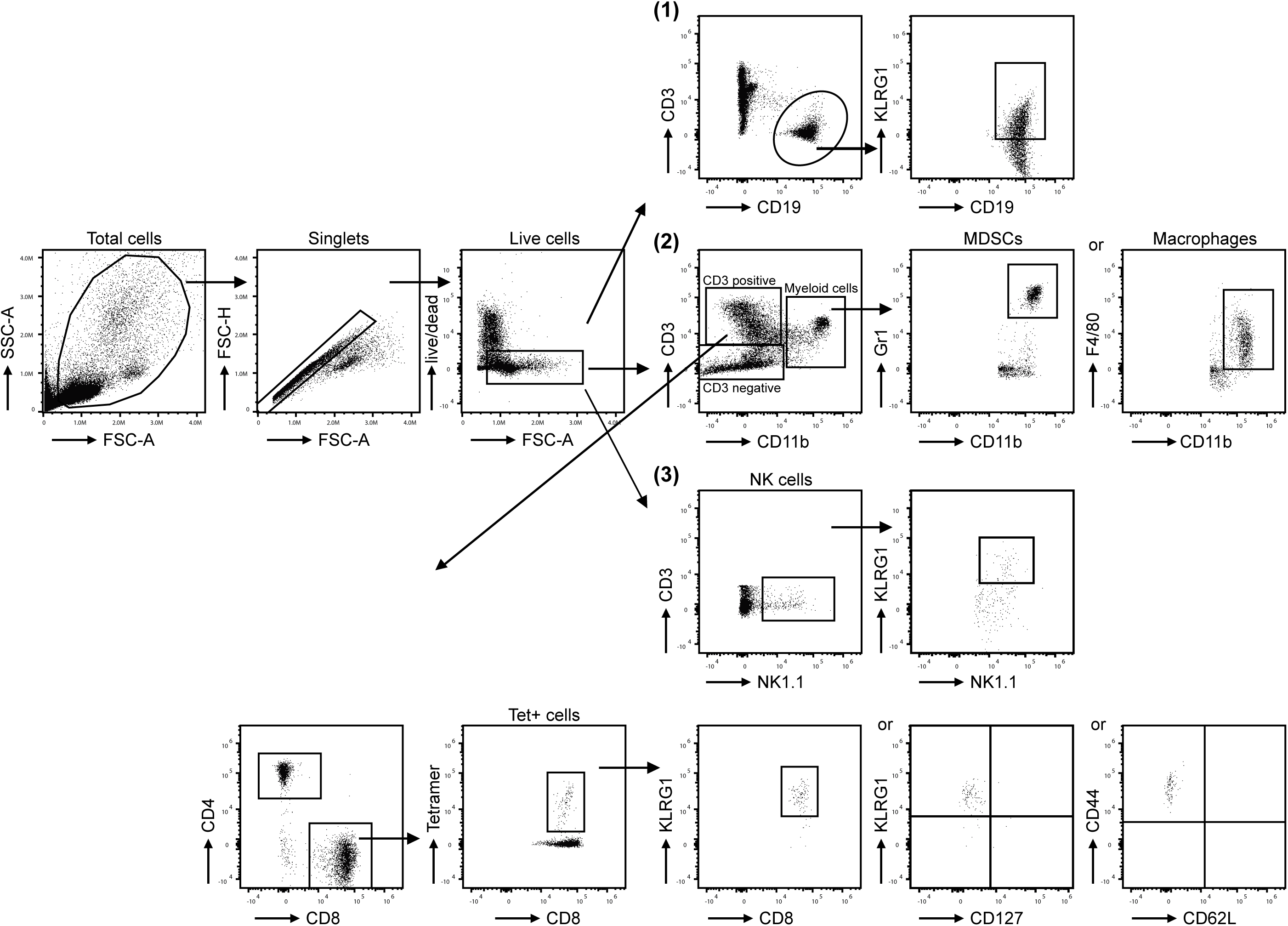
Gating strategy used for analysis of immune cell subsets. Representative flow cytometry gating strategy used to identify immune cell populations analyzed throughout the study. Sequential gating steps are shown, including exclusion of doublets and dead cells, followed by lineage-specific markers for T cells, B cells, NK cells, myeloid subsets, and antigen-specific CD8^+^ T cells using MHC-I tetramers.

**Figure S2.**
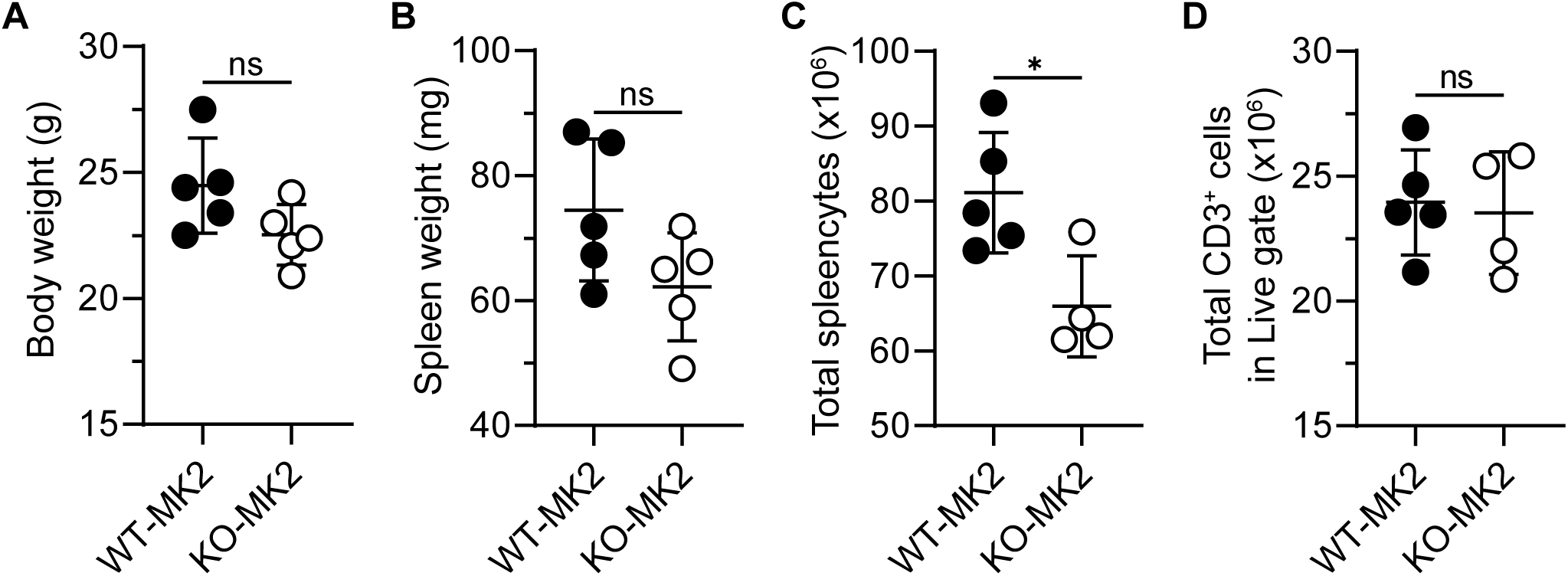
MK2 deficiency does not impact baseline body weight or lymphoid tissue composition in naïve mice. **A.** Body weights (g) of age-matched naïve WT and KO-MK2 male and female mice. **B.** Spleen weights (mg) from naïve WT and KO-MK2 male and female mice. **C.** Total splenocyte counts in naïve WT and KO-MK2 mice. **D.** Total numbers of CD3^+^ T cells in spleens of naïve WT and KO-MK2 mice (n = 4-5 per group). All data are presented as mean ± SD. Statistical analysis was performed using an unpaired two-tailed Student’s t test. Significance: *, P < 0.05; ns,not significant (P > 0.05).

**Figure S3.**
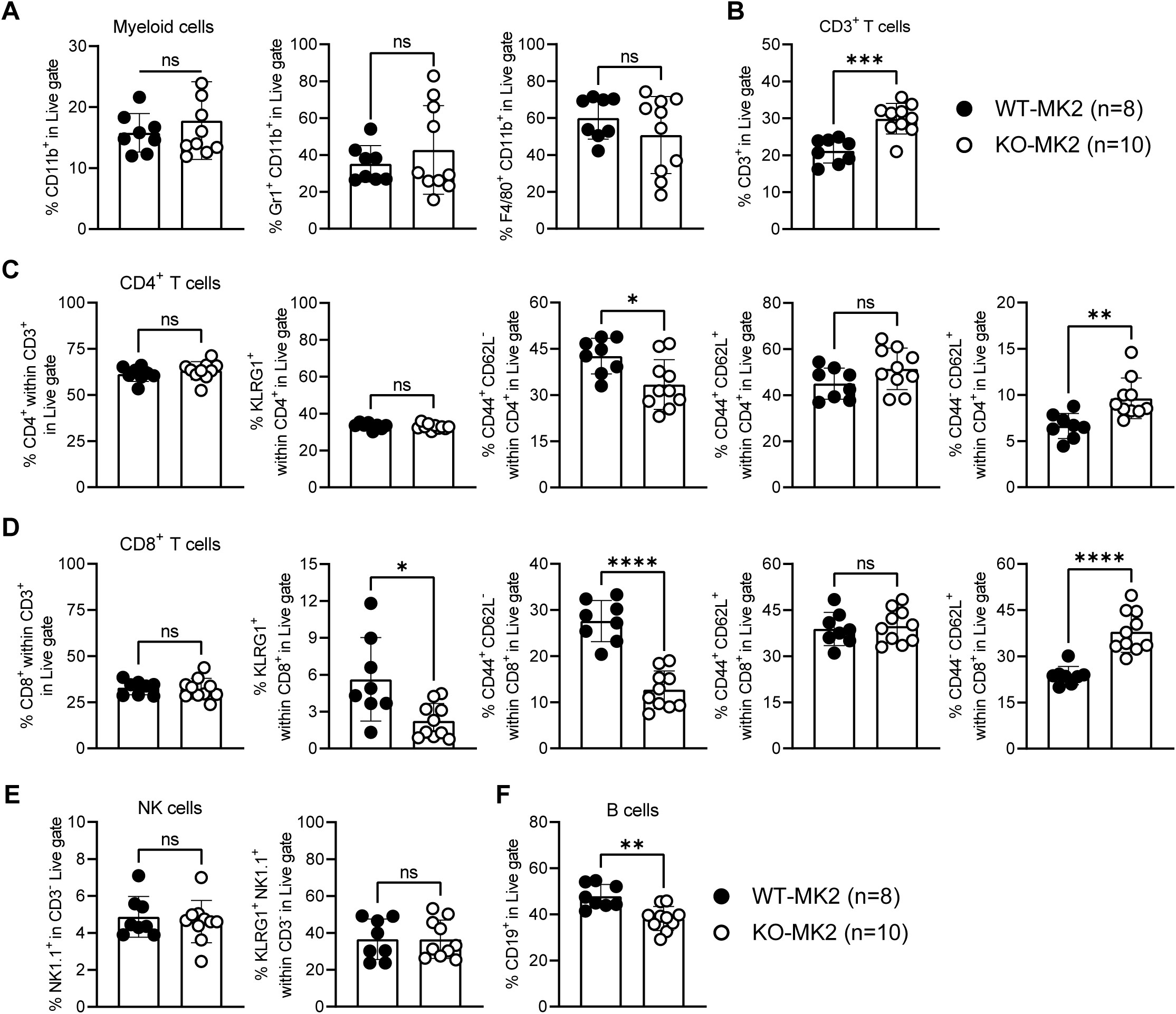
Increased CD3^+^ T-cells and naive CD8^+^ T-cells, and reduced CD4^+^ and CD8^+^ effector-memory T-cells and CD19^+^ B-cells in the peripheral blood of MK2-deficient mice. **A.** Myeloid cells, **B.** CD3^+^ T cells, **C.** CD4^+^ T cells, **D.** CD8^+^ T cells, and **E.** NK cell subsets in peripheral blood from naïve WT and MK2-KO male and female mice were measured by flow cytometry using an extension of the gating strategy shown in Figure S1. Statistical analysis was performed using an unpaired two-tailed Student’s t test. Significance: *, P < 0.05; **, P<0.01; ***, P < 0.001; ****, P < 0.0001; ns, not significant (P > 0.05).

**Figure S4:**
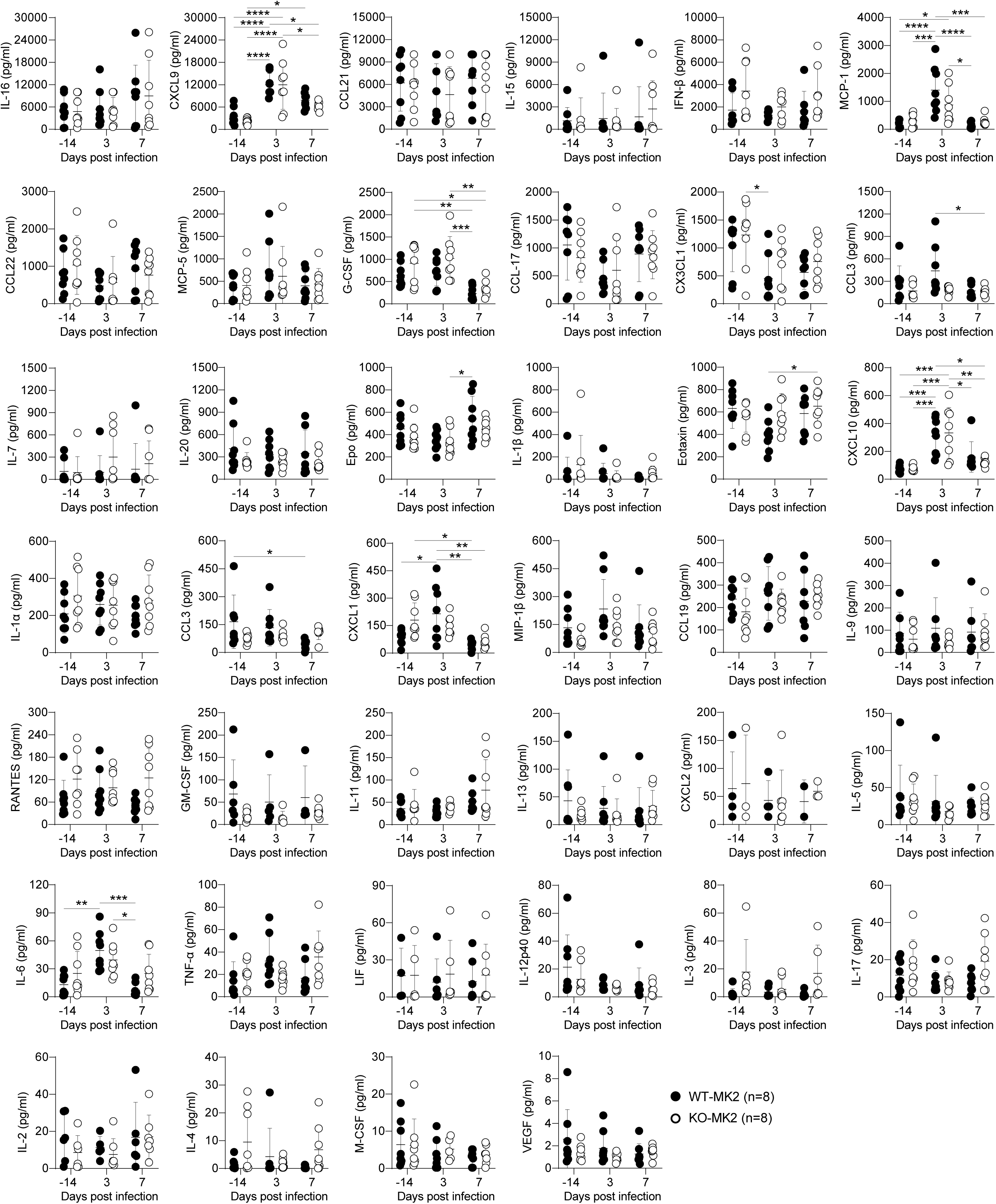
Systemic cytokine responses in MK2-deficient mice. Levels of cytokines and chemokines in the serum of WT and KO-MK2 mice were measured at baseline (day –14) and on days 3 and 7 post-MCMV infection. Cytokine concentrations are shown in pg/mL. All samples were run in duplicate, and the mean value is shown (n = 8 mice per group). Each dot represents an individual mouse. Data are presented as mean ± SD. Analytes are ordered from top to bottom based on the magnitude of their systemic induction. Statistical comparisons were performed using two-way ANOVA followed by Tukey’s multiple comparisons test. Significance is indicated as follows: *, *P* < 0.05; **, *P* < 0.01; ***, *P* < 0.001; ****, *P* < 0.0001; ns, not significant.

## Notes

### Competing Interest Statement

The authors have declared no competing interest.

